# Flexible foraging effort shapes Adélie penguin reproductive success across Antarctica and sea ice conditions

**DOI:** 10.1101/2025.07.05.663228

**Authors:** Téo Barracho, Gaël Bardon, Aymeric Houstin, Michaël Beaulieu, Pierrick Blanchard, Robin Cristofari, Lana Lenourry, Thierry Raclot, Clarisse Tabillon, Daniel P. Zitterbart, Nicolas Lecomte, Céline Le Bohec

## Abstract

Understanding how species deal with diverse environmental conditions is crucial for predicting their future responses to climate change. In Antarctica, changing sea ice dynamics threaten ice-dependent species. Whilebehavioural adjustmentscould help mitigate these threats, their effectiveness under future environmental conditions and potential thresholds remain unclear. Long-term studies and circumpolar approaches could address this challenge. Our 15-year study (2010-2024) of over 23,000 foraging trips from Adélie penguins (*Pygoscelis adeliae*) in East Antarctica, identified a non-linear relationship between sea ice concentration (SIC) and foraging effort when landfast ice was absent, a situation anticipated to define the species’ future habitat during chick-rearing. Minimal effort occurred at 10-20% SIC, but more than doubled with the presence of landfast ice or heavy pack-ice, with females showing greater sensitivity to these ice changes. Acircumpolar analysis of similar data demonstrated that foraging effort during chick-guarding predicted reproductive success across ten Adélie penguin populations. Reproductive success remained high until foraging trips exceeded approximately 29 hours, after which performance declined sharply. This indicates that while behavioural flexibility can buffer against moderate environmental changes, it has clear limits. Our findings establish foraging trip duration as a valuable early warning indicator for population responses to environmental change.

## Introduction

Global change is exerting an increasing pressure on ecosystems, challenging their resilience, and threatening species with extinction^1,2^. As changes in species abundance stem from changes in vital rates such as survival and reproduction^3^, adopting a demographic framework is a powerful way towards understanding and forecasting the impacts of global change on biodiversity^4–7^. For example, widespread declines in bird offspring production^8^ may foreshadow future population declines^9^.

Nevertheless, such declines are not homogeneous, and some species are able to withstand, or even benefit from, the modifications in their environment associated with global change^10,11^. While compensation among vital rates (e.g., increases in one vital rate proportional to decreases in another^12,13^ may explain part of this resilience, a major and rapid mechanism through which species can respond to changing environments is by modifying their behaviour^14–17^. Behavioural modifications observed at the population-level in response to changing environments (adjustments) can arise through a variety of processes. These involve reversible changes within the same individuals (contextual plasticity), or a turnover of individuals with similar (developmental plasticity) or distinct (genetic evolution) genotypes^18–21^. Regardless of the origins of behavioural adjustments (here defined at the population level), species or population are often able to adjust key behaviours such as breeding phenology or activity budgets, which can help them mitigate or even entirely escape the demographic consequences of mismatched food resources^22–25^, unpredictable food shortages^26–28^ or rising temperatures^29,30^.

While behaviour is considered an early response to stressors^9^, the limitations and potential thresholds of behavioural adjustments in allowing species to cope with changing environments are poorly known^14^. Specifically, examples charting the efficacy of behavioural adjustments for buffering demography declines with increasing environmental constraint are few^31,32^. Shorter-term studies can provide a binary responseto the efficacy of behavioural adjustments (e.g., sufficient: ^27,33–35^ ; insufficient: ^36,37^). However, a limited cover of environmental conditions will hide the full suiteof possibleresponsesand flexibility^38^. In this context, long term studies and cross-populations syntheses (e.g., space for time substitutions^39^) can help resolve this challenge^40,41^.

Antarctica presents an ideal natural laboratory for exploring these questions, with rising temperatures, shifts in wind and precipitation patterns, changing sea ice dynamics, and fisheries development all combining to affect food webs across the continent^42–49^. Many Antarctic species are long lived and as such are likely to rely heavily on plasticity (physiological or behavioural) for responding to these changes^43,50^. Among them, the Adélie penguin (*Pygoscelis adeliae*) is the focus of several long-term research programs around the continent^51–56^ making this species suitable for continent-wide approaches.

Adélie penguins are reliant on sea ice^51,57,58^, a dominant feature of the Southern Ocean^59,60^. Sea ice can impact Adélie penguin food availability in two ways: it controls the population dynamics (abundance) of its main prey species (i.e. Antarctic and ice krill, Antarctic silverfish, *Euphausia superba & E. crystallorophias, Pleuragramma antarctica*, respectively^51,61,62^, and can also constrain penguins’ accessibility to these prey. While non-breeding Adélie penguins can escape local food constraints by travelling thousands of kilometers^63,64^, they are central place foragers bound to their breeding site during the breeding season^65^.

During the chick-guard stage of the breeding cycle, foraging penguins are particularly exposed to local changes in food availability, with the regular provisioning of young and thermally dependent chicks requiring them to forage over smaller areas compared to other breeding and non-breeding stages ^64,66,67^. The availability of food during chick-guard can be affected by sea ice at different timescales, the main of which appears to differ across Antarctic regions. In areas where sea ice persists near colonies during chick-guard, such as the Ross Sea or East Antarctica, concurrent sea ice conditions are considered a reliable predictor of local food availability ^56,68–71^. However, in areas where sea ice seldom persists after incubation (e.g., the western Antarctic Peninsula^63,72^), variations in chick-guard food availability are thought to be mostly driven by sea ice conditions prior to chick-rearing, such as in the previous winter or in early spring^52,73^.

Regardless of its drivers, chick-guarding penguins can respond to variable food availability by adjusting their foraging effort, for example by performing longer trips when food availability is reduced ^56,74,75^. While such adjustments may lead to higher energy expenditure^74^, they could allow foraging parents to find sufficient food to sustain their chicks. Nevertheless, the extent to which foraging adjustments allow penguins to buffer reproductive success against variable food availability is poorly known ^76^. For populations where chick-guard food availability is driven by concurrent sea ice conditions, the type of ice encountered by penguins likely affects this outcome^57,76^. For instance, the persistence of landfast ice (consolidated sea ice anchored to the coastline) in front of breeding colonies during chick-guard leads to much longer foraging trips because penguins have to travel by walking rather than swimming^56,77^. In such cases, reproductive success is often severely reduced ^57,78^. In the absence of landfast ice, foraging penguins can still face varying concentrations of pack ice, ranging from open water (0% Sea Ice Concentration, SIC) to heavy pack ice (up to 80% SIC). In this scenario, the foraging response of penguins and the consequences on reproductive success are not fully resolved, although foraging effort was found to be minimal at intermediate 10-20% SIC when measured over a narrow range of SIC (0-30% SIC)^69,71^. Because most Adélie penguin populations typically have direct access to open water during chick-rearing (i.e. with an absence of landfast ice)^51,79^, and because landfast ice is expected to recede further under climate change^59^, understanding penguin responses to the variable pack ice conditions they predominantly encounter is crucial for projecting their demographic responses to future environmental changes.

Here, we combine a unique 15-year-long time series of Adélie penguin’s foraging effort (trip duration) and reproductive success from Adélie Land, Antarctica, with an additional circumpolar analysis of previously published data spanning the species range to address two main questions. First, we ask how Adélie penguins at our Adélie Land study site adjust their foraging effort to variable food availability, as proxied by sea ice conditions in their chick-guard foraging area. Specifically, we investigate how penguins respond to a wide range of sea ice concentrations (0 -99%) while contrasting both ice types (landfast vs. pack ice) to match the expected icescape the species will face in the coming decades. Second, we explore the consequences of these foraging effort adjustments on reproductive success. We predict that increased foraging effort, as reflected by longer foraging trips, will lead to lower reproductive success^56,76^, even in the absence of landfast ice. By scaling-up the analysis to include other colonies, we aim at quantifying the relationship between foraging effort and reproductive success across the species distribution. Such data integration over a wide range of conditions allows us to understand the relevance of foraging effort as a fundamental mechanism linking environmental variability to reproductive success, providing a robust baseline for projecting the species responses to changing sea ice dynamics.

## Results

Our 15-year dataset (2010-2024) of Adélie penguin foraging trip duration during the chick-guard period (inter-annual average: Dec. 23^rd^ to Jan. 19^th^) encompassed 23,459 trips from 565 individual penguins that performed a total of 1,779 breeding cycles. On average, each individual was included in our dataset in 3 different breeding seasons (range: 1-12). The annual number of trips averaged 782 trips per year-sex combination (range: 10-3,013, Table S1). Landfast ice persisted in front of the colony through the chick-guard stage in five years (2012, 2014, 2015, 2017, 2018), when minimal distances to open water ranged from 4 km in 2012 to 60 km in 2017. For these five years, foraging penguins faced elevated sea ice concentrations (SIC) ranging from 43 to 99% (mean = 58 ± 0.33%; Fig. 1). For all the other years, landfast ice was absent in front of the colony during the chick-guard stage and penguins experienced a lower average (20 ± 0.14%) but a wider range (0-84%) of SIC, which even included near-total open water conditions (Fig. 1).

**Figure 1:**
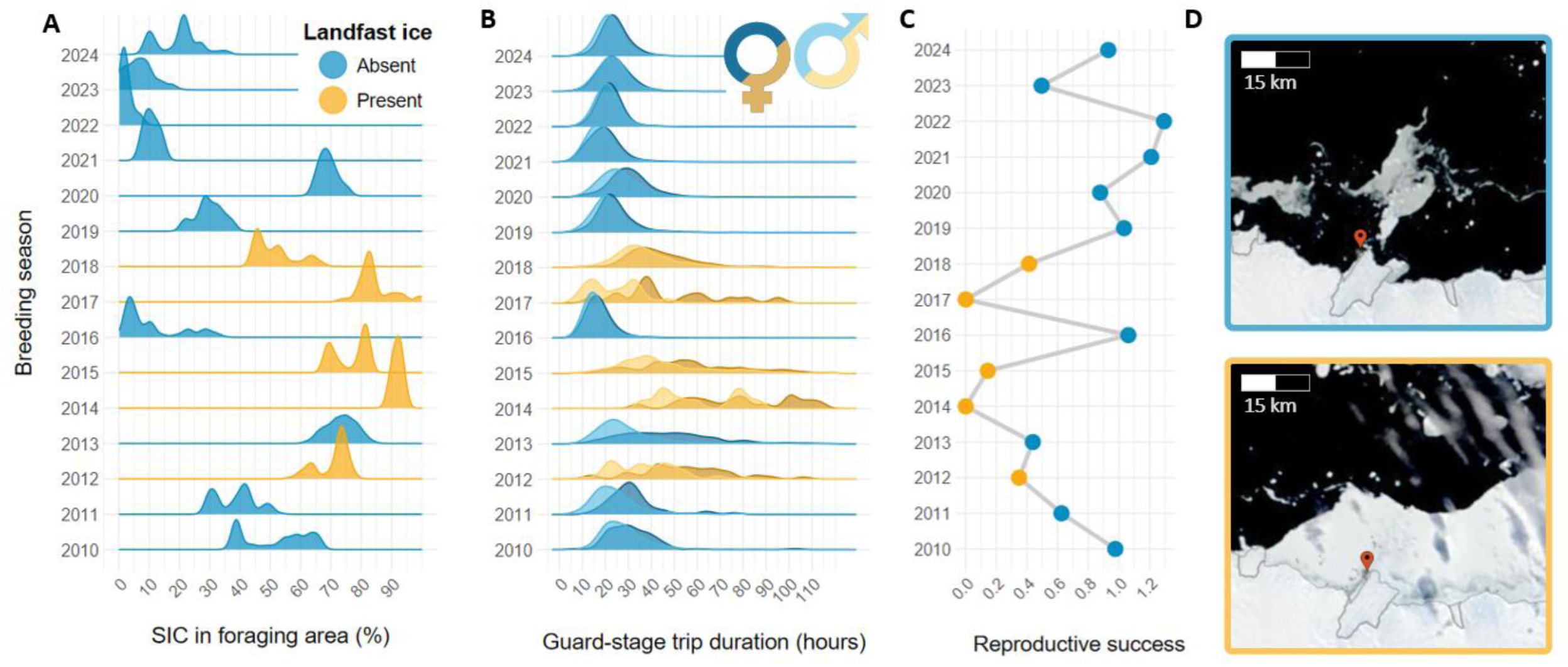
Cascading relationship between sea ice concentration (SIC), foraging trip duration and reproductive success for Adélie penguins (*Pygoscelis adeliae*) breeding in Pointe Géologie archipelago, Adélie Land, East Antarctica over the 2010-2024 period. Two contrasting scenarios of access to open water during chick-rearing are apparent: direct access to open water is either possible (*landfast ice absent*, blue) or precluded by the presence of solid landfast ice in front of the breeding colony (*landfast ice present*, yellow). **A)** Annual density distribution of SIC experienced by Adélie penguins during their foraging trips throughout the chick-guard stage. **B)** Annual density distribution of foraging trip duration (n = 23,459 trips) for chick-guarding male and female Adélie penguins from a ca. 250 pairs breeding patch (study colony) located on Île des Pétrels, Pointe Géologie archipelago. **C)** Reproductive success time series for the study colony (2010-2024 average ± SD = 0.66 ± 0.43 chicks per breeding pair). Reproductive success was calculated as the number of chicks fledged per breeding pair and can exceed 1 because Adélie penguins can raise two chicks^51^. **D)** Satellite imagery representative of the absence (top) and presence (bottom) of landfast ice in front of the study colony (red marker). MODIS satellite imagery was downloaded from the NASA Worldview application (https://worldview.earthdata.nasa.gov), part of the NASA Earth Observing System Data and Information Sy stem (EOSDIS, top and bottom images from Jan 10th, 2022, and Jan 1st, 2018, respectively).

### Trip duration response to the sea ice scape

In line with this sea ice variability, foraging trip duration during the chick-guard period also varied considerably (mean = 23.4 ± 0.08 h). The presence of landfast ice in front of the breeding colony was the main driver of foraging trip duration. Trips were twice as long in years of landfast ice presence(mean = 43.9 ± 0.46, n = 1,621 trips) compared to when landfast ice was absent (mean = 21.9 ± 0.06 h, n = 21,838 trips, Welch’s t-test p < 0.001). Over the range of SIC values shared by both landfast ice presence/absence scenarios (i.e. 43-84%, Fig. 2), trips were, on average, 1.4 times longer when landfast ice was present than when it was absent (Welch’s t-test p < 0.001). In the absence of landfast ice, we found strong support for a non-linear effect of SIC on trip duration (GLMM vs. GAMM ΔAIC = 450; Table S2; Fig. 2), with the model-predicted 10% shortest trips (range: 19-20 h) occurring within 8-18% SIC (Fig. 2). Foraging trip duration lengthened rapidly between 20 and 40% SIC, but increased more slowly at ca. 70-80% SIC, with the model-predicted 10% longest trip duration (range: 29-37 h) occurring above 76% SIC. Comparison of raw data at optimal (8-18%) versus high SIC (>76%) indicates that this represents an average 2.5-fold increase in trip duration. At the other end of the SIC range (<8% SIC), we also observed an increase in trip duration, but of much lower magnitude (1.04 -fold increase compared to optimal 8-18% SIC; Fig. 2). Similar to the landfast ice absent scenario, we found that foraging trip duration lengthened with increasing SIC when landfast ice was present.

**Figure 2:**
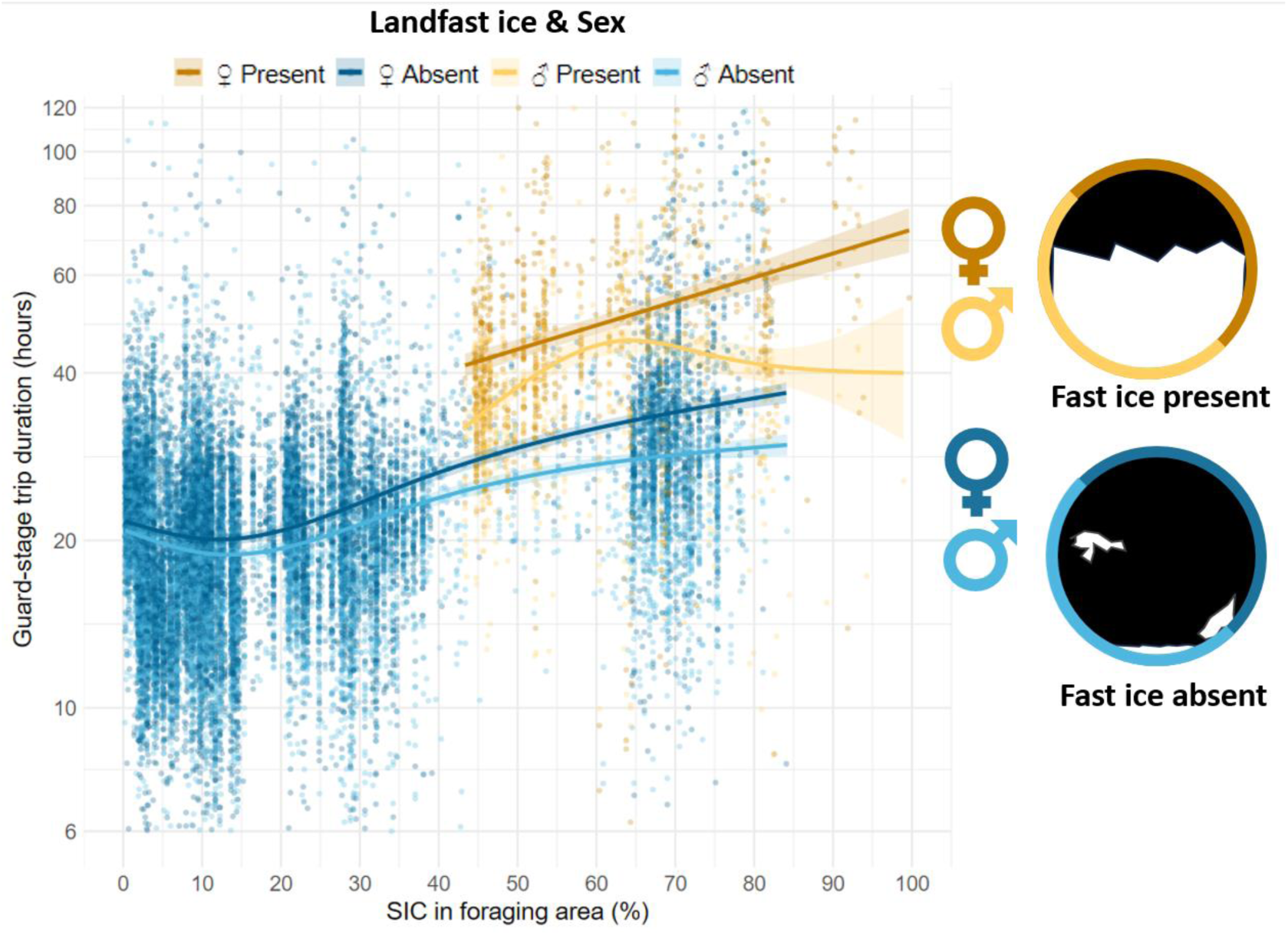
Relationship between the foraging trip duration (log scale) of chick-guarding Adélie penguins (*Pygoscelis adeliae*) and sea ice concentrations (SIC) for two contrasting scenarios of access to open water: direct access to open water is either possible (*landfast ice absent*, blue) or precluded by the presence of solid landfast ice in front of the breeding colony (*landfast ice present*, yellow). Trip duration data (n = 23,459 trips) was collected using a RadioFrequency Identification System deployed around a breeding patch of ca. 250 pairs within the Pointe Géologie archipelago population (Adélie Land, Antarctica). Lines and asso ciated confidence intervals represent the predictions of the Generalized Additive Mixed Models (GAMM) examined in Table S2. When landfast ice is absent, the 10% lowest trip durations predicted by the GAMM are attained for SIC ranging 8-18% (raw data trip duration mean ± SE = 19.4 ± 0.09 h). Conversely, the predicted 10% longest trip durations occur at high SIC (>76%, raw data trip duration mean ± SE: 47.5 ± 2.17 h), indicating an average 2.5 -fold increase between optimal and high SIC. When direct access to open water is precluded by the presence of landfast ice in front of the breeding colony, trip durations are lengthened, averaging 43.9 ± 0.46 h (raw data). This represents an average 1.4-fold increase in trip duration compared to landfast ice absence conditions over the comparable range of SIC (43-84%), and an average 2.2-fold increase compared to the optimal 8-18% SIC in the absence of landfast ice.

### The relationship between trip duration and SIC is mediated by sex

Chick-guard foraging trip durations differed between male and female Adélie penguins in both landfast ice scenarios (Table S2, Fig. 1B). Overall, females exhibited foraging trips almost 2 hours longer on average compared to males (females: mean = 24.4 ± 0.11 h, n = 12,135 trips *vs*. males: mean = 22.4 ± 0.10 h, n = 11,324 trips, Welch’s t-test p < 0.001). Regardless of the landfast ice scenario, sex-differences were exacerbated under increasing SIC, as shown by the overall relationship between SIC and the annual male-female trip duration difference (LM, β = 0.22 ± 0.03, p < 0.001, R^2^ = 0.79; Fig. S2). Specifically, while the difference in trip duration between sexes was under 2 hours below 20% SIC, it increased beyond 14 hours above 70% SIC (Fig. S2). This relationship was maintained when landfast ice was absent (LM, β = 0.15 ± 0.04, p = 0.002, R^2^ = 0.70; Fig. S2).

### Consequences on reproduction

Our 15-year study encompassed substantial interannual variation in reproductive success, ranging from 0 (2014, 2017) to a maximum of 1.29 chicks/pair (2022), with an overall mean of 0.66 ± 0.43 chicks/pair. Reproductive success was negatively related to mean SIC in the chick-guard foraging area (Tweedie GLM: β = -0.020 ± 0.005, p < 0.001, pseudo-R^2^ = 0.58, Fig. 3A) and was lower in years when landfast ice was present in front of the colony (Welch’s t-test, p < 0.001, Fig. 1C). We found that longer foraging trips during chick-guard were associated with a linear reduction in reproductive success, with a model-predicted 4.3% decrease in reproductive success every 1 -h increase in mean trip duration (Tweedie GLM: β = -0.044 ± 0.006, p < 0.001, pseudo-R^2^ = 0.73, Fig. 3B). This relationship was maintained when considering only years without landfast ice (Tweedie GLM: β = -0.035 ± 0.007, p < 0.001, pseudo-R^2^ = 0.48). Overall, the mean trip duration of females was a better predictor of reproductive success (pseudo-R^2^ = 0.78) than that of males (pseudo-R^2^ = 0.57), including in years when landfast ice was absent (Table S3).

**Figure 3:**
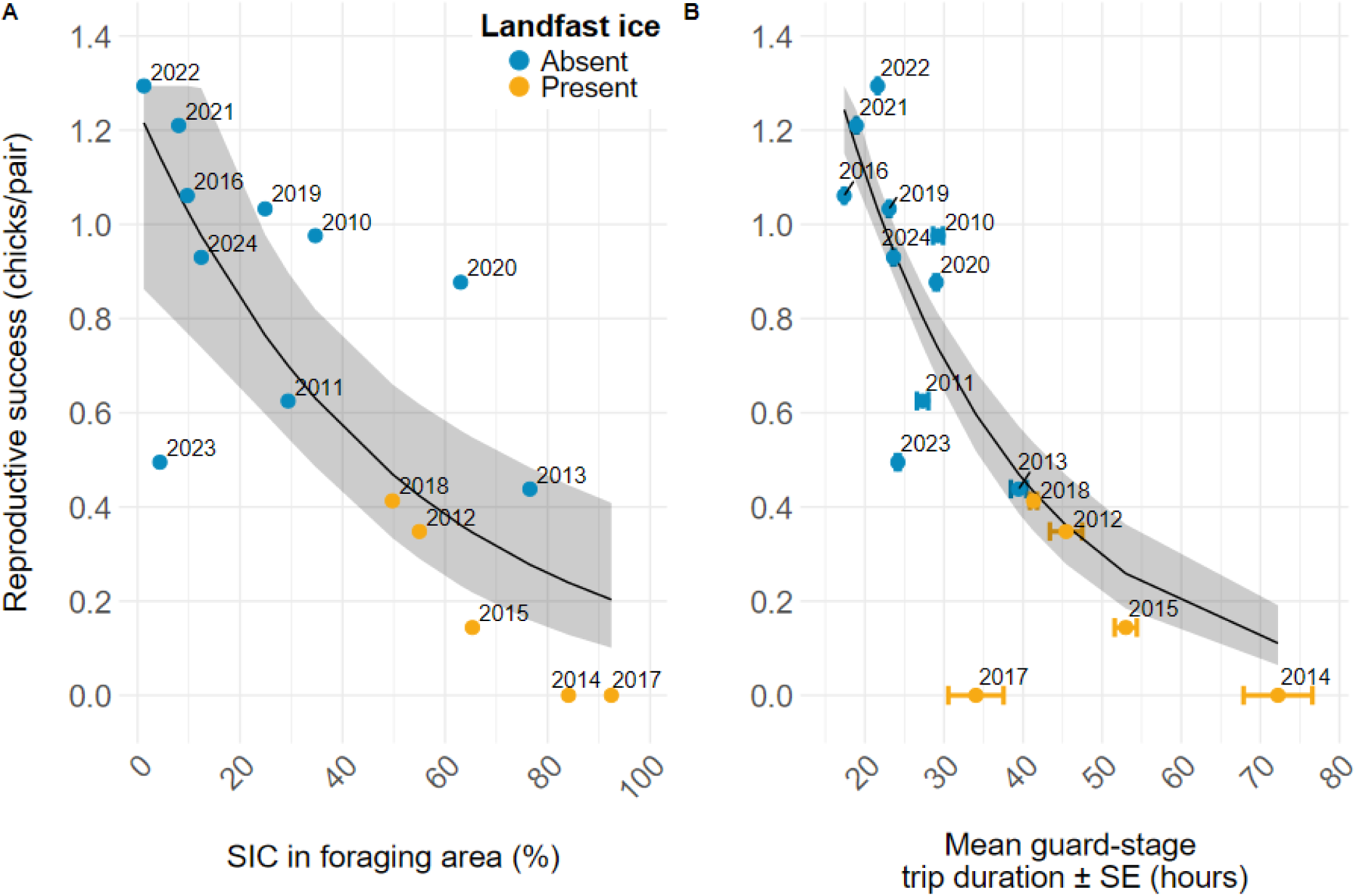
Relationship between Adélie penguin (*Pygoscelis adeliae*) reproductive success (chicks/pair) and A) sea ice concentration (SIC) in the foraging area used during chick-guard (pseudo-R^2^ = 0.58); and B) mean foraging trip duration (pseudo-R^2^ = 0.71) of chick-guarding Adélie penguins breeding on Île des Pétrels, Pointe Géologie archipelago, Adélie Land, Antarctica (2010-2024). Black lines and associated confidence intervals are from generalized linear models with a Tweedie distribution. When only years without landfast ice are considered (blue), the relationship between SIC and reproductive success (A) is not statistically significant (p = 0.058, pseudo-R^2^ = 0.25), while the relationship between mean trip duration and reproductive success (B) remains significant (p = 0.003, pseudo-R^2^ = 0.46).

The circumpolar analysis including data from 9 other Adélie penguin populations spread across the species range (see supplementary text) yielded an additional 49 concurrent measures of trip duration and reproductive success. Over the total 64 observations, covering East Antarctica, the Ross Sea, and the Western Antarctic Peninsula (Fig. 4), reproductivesuccessaveraged 0.88 ± 0.37 chicks/pair, while mean trip duration averaged 29.7 h (range 9.6-72.2 h). Similar to our study site, we found a significant relationship between reproductive success and trip duration (Table S4, p < 0.001, marginal R^2^ = 0.74), but with non-linear effects of trip duration on reproductive success (linear vs. quadratic GLMM ΔAIC = 33, Table S4). Model-predicted reproductive success remained well above average (>1 chicks/pair) below approximately 30 h of trip duration, before declining more sharply (Fig. 4). To investigate this potential threshold, we carried-out a piecewise regression analysis *a posteriori* and estimated the breakpoint at 29.2 h (bootstrapped 95% confidence interval: 17-40 h, Fig. S3).

**Figure 4:**
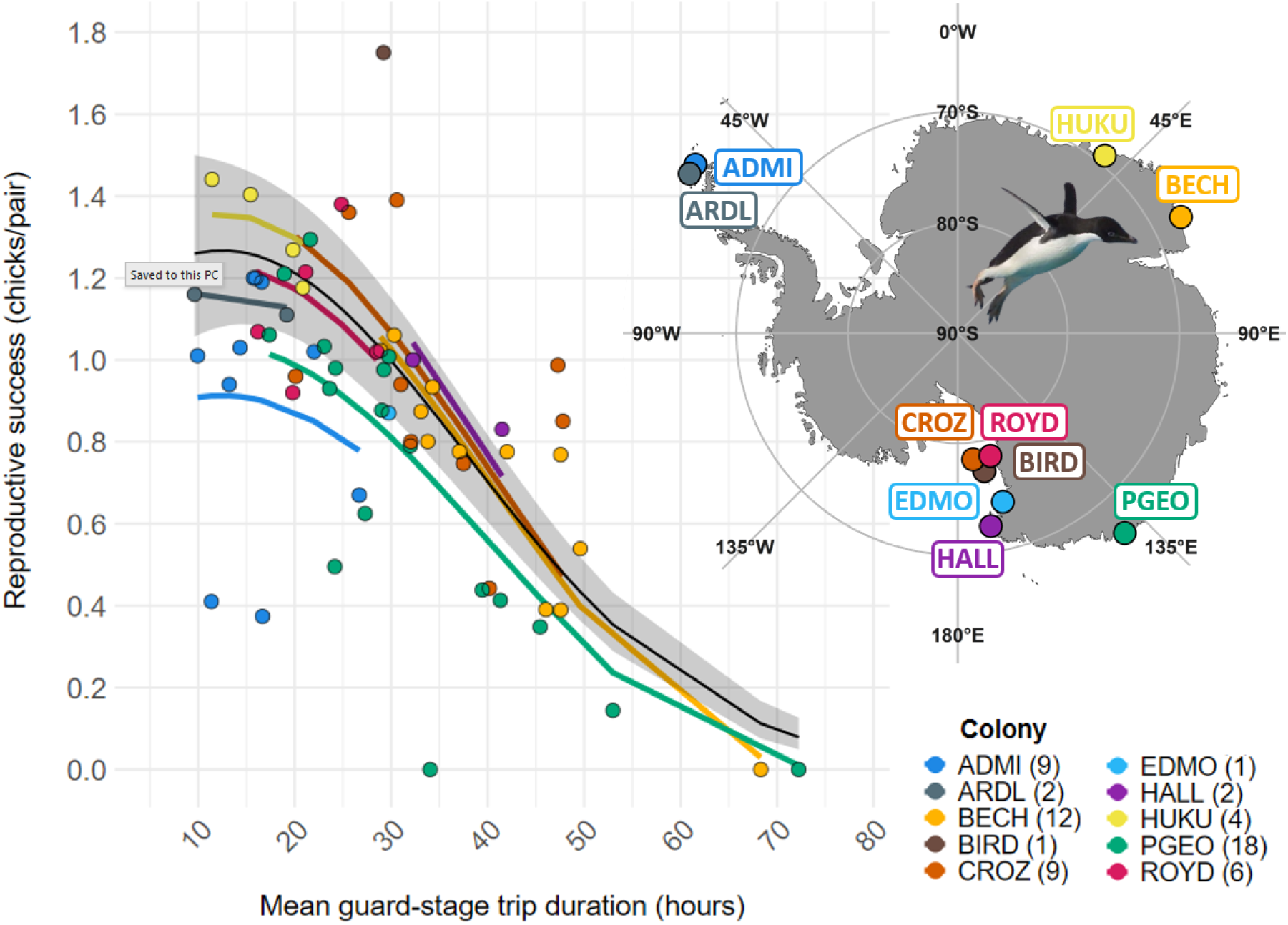
Relationship between Adélie penguin (*Pygoscelisadeliae*) reproductivesuccessand mean foraging trip duration across 10 coloniesaround Antarctica. Theregression lines and associated confidence interval for the general relationship (black line) are from a Generalized Linear Mixed Model (GLMM) including colony as a random effect (marginal R^2^ = 0.74). Numbers in parentheses after colony abbreviations are colony-specific sample sizes (number of breeding seasons with concurrent reproductive success and trip duration data). Data for Pointe Géologie mainly comes from the present study (n = 15 seasons) but also includes data retrieved from the literature^84,96,169^. Data for all other colonies was retrieved from the literature, and full references and raw data can be found in supplementary text. Abbreviations for colony names: ARDL, Ardley Island^85^; ADMI, Admiralty Bay^49,170^; HUKU, Hukuro Cove^56^; BECH, Béchervaise Island^76^; PGEO, Pointe Géologie archipelago; HALL, Cape Hallett^148^; EDMO, Edmonson Point^55,86^; ROYD, Cape Royds^68,161,171,172^; BIRD, Cape Bird^161^; CROZ, Cape Crozier^68,111,161^.

## Discussion

Behavioural adjustments to fluctuating environments are likely to be a central mechanism through which species can respond to climate change and other human pressures ^17,80^. As such, capturing these behavioural adjustments over their largest possible extent and understanding their effectiveness for buffering demographic rates against environmental variability is crucial for identifying potential perturbation thresholds^81^ and species tipping points^82,83^, as well as for designing reactive conservation policies^9,14^. Here, we showed how Adélie penguins responded to an unprecedentedly wide range of sea ice variability through flexibleforaging effort andidentified a non-linear linkage between foraging effort and reproductivesuccess acrossthe species pan-Antarcticrange. Our results provide a powerful template to assess Adélie penguins’ response to anticipated changes in Antarctica.

### The sea ice scape constrains foraging effort

The strong, positive relationship between landfast ice presence (sea ice anchored to the coastline) and trip duration found in our 15-year study corroborates previous findings^69,76,78^, confirming that the need to walk over ice rather than swim significantly increases foraging effort^56^. Importantly, a remarkable contribution of our long term study is the characterization of the species response over a wide spectrum of sea ice concentration (SIC) in the absence of landfast ice, a condition that predominates for most Adélie penguin populations^51,79^ and is likely to become increasingly common as landfast ice recedes over the coming century^59^.

The non-linear relationship we observed between foraging effort and SIC (Fig. 2) aligns with previous studies in the Ross Sea^69,77^ and East Antarctica^71^ showing optimal foraging at intermediate sea ice concentrations (8-18% SIC). However, by extending the range of SIC covered from 0-30% (ref.^69^) to 0-84%, our study revealed that foraging effort increases substantially at high SIC (>76%), even in the absence of landfast ice. Although such high SIC values are rarely attained in the absence of landfast ice (Fig. 2), dense pack-ice still presents a significant challenge to foraging penguins, with trip duration increased up to 2.5-fold compared to optimal conditions. This value is comparable with the average 2.2-fold increase in trip duration between optimal conditions (8-18%SIC, no landfast ice) and landfast ice conditions.

Interestingly, male and female Adélie penguins differed in their response to challenging sea ice conditions. While females consistently undertook longer trips than males, matching previous observations locally^67,84^ and at other colonies around Antarctica^76,85,86^, we showed that females experienced a steeper increase in trip duration with increasing SIC compared to males (Fig. S2). Such differences could be related to sex-specific diets, with males and females preferentially targeting fish and krill, respectively^84,86^, or to differences in body condition at the time of chick-guard, with females in poorer condition more proneto engagein longer self-feeding trips. Sex-specific sensitivity to foraging conditions could also signal the potential for sex specific threats (e.g., ref.^87,88^), although no difference in survival were found between male and female breeders in the Ross Sea ^89^.

### How does sea ice modulates foraging effort?

The increased effort at both very low (0% SIC) and high (>50% SIC) concentrations likely reflects different limitations on foraging efficiency. In open water (0% SIC), Adélie penguins may struggle to locate and exploit prey patches efficiently^70,90^. As Antarctic krill and silverfish aggregate near sea ice edges^61,91^, the complete absence of ice could remove these aggregating structures. The absence of sea ice during summer may also directly reduce the local abundance of prey species associated with sea ice year-round (e.g., *E. crystallorophias*), although the abundance of both *E. crystallorophias and E. superba* has been more convincingly linked to winter ice cover rather than summer SIC^62,92,93^. At the opposite end of the SIC spectrum, high ice concentrations physically impede access to prey, thereby reducing food accessibility through increased travelling costs^56,76^. The optimal intermediate SIC (10-20%) therefore likely provides sufficient ice-edge habitat for prey aggregation, while maintaining adequate open water access for efficient travel and foraging ^71^.

### Foraging within the future sea ice scape

Another keyresult is that, over the course of our study, very low SIC (<8%) did not significantly increase Adélie penguin foraging effort, which remained within 10% of their lowest observed effort (Fig. 2). Together with even shorter trip durations sometimes observed in the summer-ice-free western Antarctic peninsula(i.e. around 10 h)^49,85^, this suggest that Adélie penguins are able to obtain sufficient food when SIC is severely reduced within their chick-rearing foraging area. In the context of ongoing and projected sea ice loss^44,45,59,94^, this is a crucial finding for understanding and anticipating the species’ response under expected environmental shifts. Nonetheless, expected sea ice declines are not confined to the summer season, and will also likely affect winter ice dynamics. Because winter sea ice is a critical habitat and driver of krill abundance during summer^62,93,95^, the ability of Adélie penguins to mitigate lower ice concentrations during summer may not prove to be a sufficient response if waning winter sea ice further alters summer food abundance. Such lags could explain unexpected observations of lower reproductive success despite low summer SIC values at our study site ^96^ or in the western Antarctic Peninsula (Fig. 4), but further concurrent data on food abundance, foraging effort and reproductive success in low SIC contexts will be key to test this hypothesis.

While counter-intuitive, predicted increase in air and water temperatures in Antarctica could also foster local increases in landfast ice and dense pack-ice persistence during the breeding season, which we both showed to significantly impact Adélie penguin foraging effort. Grounded icebergs can indeed act as anchor points for landfast ice, and glacier calvings are expected to increase under climate change-driven changes in temperatures, windstorms, and rain regimes^59^. While the location and duration of iceberg grounding events remain highly unpredictable^59^, such mechanisms could explain the rising landfast ice trends in certain regions, notably East Antarctica^97^.

### Foraging effort is a reliable predictor of reproductive success

A major outcome of our study is the robust relationship between foraging trip duration during chick-guard and reproductive success, both at our study site (Fig. 3B) and across the species’ range (Fig. 4). This relationship persisted even when considering only years without landfast ice, demonstrating that varying pack-ice concentrations alone can significantly impact reproductive outcomes. While substantial variability remains, the non-linear relationship we revealed over different Antarctic regions—from the ice-heavy Ross Sea to the increasingly maritime West Antarctic Peninsula— highlights the limits of flexible foraging effort as a mechanism for coping with changes in food availability during a critical phase of the species’ breeding cycle.

### Why is chick-guard trip duration a good predictor of reproductive success?

Reproductive failure often peaks during early chick-rearing in Adélie penguins^98–100^ as well as other seabirds (e.g., refs.^101–103^ but see ref.^104^). This makes the chick-guard period, and the ability of breeders to provision their chicks with food at a high rate by performing short trips, critical for reproductive success. While trip duration is constrained by foraging conditions encountered during chick-rearing, it also depends on the conditions encountered during incubation. Adélie penguins indeed use incubation foraging trips to regain the body condition that they subsequently lose during chick-guard, in order to maximize energy allocation towards the offspring^105,106^. Accordingly, failure to regain body conditions during incubation leads to longer foraging trips during chick-guard^69,78^. The integration of foraging conditions during both incubation and early chick-rearing thus likely makes chick-guard trip duration a reliable predictor of reproductive success.

The foraging strategy^107,108^ employed by Adélie penguins is also important in explaining this relationship. Adélie penguins operate as time-minimizers during chick-guard, with foraging parents aiming at provisioning their offspring as quickly as possible, rather than maximizing food loads ^105,109^. This is consistent with the limited stomach capacity of guard-stage chicks compared to maximal parental food loads (i.e. > 1 kg)^110^, and with observations that chicks fed more often have faster growth rates and daily survival^99,111^. Importantly, the temporal constraints on foraging parents during chick-guard are not as strong during the incubation and crèching phases, explaining failed attempts at using trip duration during incubation as an index of food availability ^112^.

### Trip duration as a tool for forecasting population reproductive performance

Seabirds are considered potent indicators of food supplies, with their behaviour more responsive to changes in food availability compared to demographic rates or abundance^113–115^. Consistently, behavioural changes sit at the forefront of the *timeline to collapse*, whereby the consequences of ramping environmental forcings should successively become observable on behaviour, morphological traits, demographic rates and ultimately abundance^9^. While this framework can improve forecasts of populations’ fate under climate change, it requires well-defined baselines against which observed changes in (behavioural) traits can be contrasted, as well as an understanding of the linkage between behaviour and demographic rates^116–119^. Here, our results provide such a template for Adélie penguins, and the existence of similar data on foraging effort and reproductive success for other central-place foragers should prompt similar endeavours. Furthermore, as the foraging effort of females appears to be a better predictor of reproductive success than that of males (perhaps because female’s longer trips can have more impact on the pair-level provisioning rate, see also^76^), our results highlight the importance of considering and including sex-specific strategies in behaviour-based assessment of population responses^120^.

Although the relationship we observed between trip duration and reproductive success was consistent across conditions and space, extra variability remains (Fig. 3B, 4) and must be considered prior to using foraging effort as a proxy for population reproductive performance. First, extreme landfast ice extent, such as that observed at our study site in 2017 (over 60 km of landfast ice between the colony and open-water) may enable chick-rearing penguins to forage only temporarily, and only for a subset of highly performant individuals^121^. Accordingly, chick-guard trip durations were particularly variable in 2017 in Pointe Géologie because some individuals were first able to forage in ice cracks, but as resources in these areas rapidly became depleted, individuals then engaged in very long trips (> 5 days, not considered in our analysis), which led to all chicks ultimately dying by mid-January^78^. Second, land-based factors independent of food availability have been shown to cause significant reductions in reproductive success for polar seabirds, including Adélie penguins^78,122–124^. Unusual weather events, particularly snow accumulation around hatching and strong winds, can indeed cause significant eggs and chick losses. Accordingly, the 2023 breeding season at our study site showed lower reproductive success than predicted by trip duration alone (Fig. 3B), largely due to heavy snowfall around hatching (personal observation). Elsewhere in Antarctica, the weather regime of the western Antarctic Peninsula is currently shifting towards a more maritime and rainy state^48^. In this region, the reproductive success of Adélie penguins appears to be particularly driven by such food-independent factors^52,123,125,126^, perhaps explaining the deviations from the general relationship between trip duration and reproductive success observed there (e.g., for Admiralty Bay, Fig. 4).

### Limits to behavioural flexibility

Identifying thresholds of behavioural flexibility beyond which environmental variability impacts demographic rates would foster our ability to gauge the scale of perturbations and forecast future population trajectories^14^. Our results, combining behavioural and demographic data from across Antarctica, suggest that the population-level threshold above which increased foraging effort ceases to buffer Adélie penguin reproductive success is about three times the minimum recorded foraging effort (i.e. around 29 h of trip duration, Fig. 3B). While a more precise breakpoint will likely be defined as more data become available, our results also suggest that behavioural flexibility does not break down immediately above this threshold. Instead, reproductive success appears to decline linearly with increasing foraging trip duration, only dropping to near zero when conditions are so unfavourable that trips exceed two times the average foraging effort (i.e. 60 h). Although this links to earlier findings that only some individuals within populations display the high foraging abilities needed for successful breeding under the harshest conditions^121,127^, it remains unclear whether such foraging effort thresholds are consistent across species — a question that future research should address. Additionally, future research should also quantify the magnitude and frequency of poor reproductive events for which population persistence could be jeopardized^128,129^.

### Behavioural adjustments could entail carry-over effects

While behavioural adjustments may entirely or partially buffer short-term fitness components (i.e. current reproduction) against environmental variability, the potential carryover effects of such adjustments could entail negative consequences on longer-term fitness components (e.g., adult survival)^130^. For example, increased foraging effort by common guillemot (*Uria aalge*) parents in response to low food availability led to decreased survival to the next breeding season ^32^. Alternately, the foraging adjustments displayed by little auks (*Alle alle*) at several colonies with contrasted resource availability allowed them not only to maintain reproductive output, but also to preserve body condition and subsequent survival^27^. In Adélie penguins, survival costs of reproduction linked to increased foraging effort have not been apparent at the population-level^131,132^. In long-lived species, we can expect the body condition threshold below which breeding individuals switch to self-feeding^105,133–135^ to be sufficiently fine tuned to prevent individuals from entering the post-breeding season with critical energy reserves. For Adélie penguins, there is considerable inter-annual variability in adult survival at some colonies^54^, and part of this variability has been linked to environmental conditions during the moult^136^. This stage of the species life cycle takes place right after reproduction^51^. There is therefore a significant opportunity for investigating carry-over effects of breeding effort on adult survival in Adélie penguins, which would provide a more complete picture into the efficacy of behavioural adjustments in buffering fitness against environmental variability.

### Conclusions and perspectives

To conclude, our study has addressed how Adélie penguins adjusted their foraging effort in response to an unprecedentedly wide range of sea ice conditions, especially in the absence of landfast ice, a feature predicted to define the future foraging and breeding habitats of Antarctic species^59,137^. Importantly, we demonstrated that chick-guard foraging trip duration, a simple measure of foraging effort, represents a valuable and readily applicable metric for monitoring the reproductive performance of a seabird’s colonies across its entire (continental) distribution, warranting its use as a potential early warning indicator of population-level declines^9^. Such ecological indicators are fundamental, and critical to the implementation of conservation and management plans in Antarctica^138^, particularly with the projected development of fisheries bolstered by sea ice losses.

## Material and methods

### Study site and monitoring design

The Adélie penguin’s breeding cycle spans the austral summer (see ref.^51^) for a general description). In Adélie Land, following arrival at the breeding colonies (October) and the laying of two eggs (November), Adélie penguin partners alternate incubation shifts and long foraging trips (5 -20 days), before switching to shorter trips upon hatching (guard stage: December-January). Broods (1 or 2 chicks) grow rapidly, and as chicks acquire self-thermoregulation abilities, parents then start to forage simultaneously to maximize food deliveries (crèche stage: January-February). Depending on the region, chicks finally fledge in January-February, with fledging mass being an important predictor of their post fledging survival^139^.

We carried out fieldwork on Île des Pétrels (66°40′S, 140°01′E), Pointe Géologie archipelago, Adélie Land, Antarctica, adjacent to the Dumont d’Urville research station. This archipelago has hosted 30,000-50,000 breeding pairs of Adélie penguins annually for the last two decades^140^, making it a medium-sized colony for this species^51,79^. From the 2006-2007 to 2023-2024 breeding seasons, we established a comprehensive monitoring system for a breeding patch comprising approximately 250 annual breeding pairs and situated within a natural canyon (hereafter study colony)^139^.

Starting in 2006-2007 (breeding seasons will be referred to by the chick-fledging year, 2007 in this case), we implanted Radio Frequency IDentification (RFID) tags in both chicks and breeding adults (n = 3,248 and n = 313 as of 2024, respectively). From the 2010 breeding season onwards, the study colony was enclosed with directional RFID gateways, strategically positioned to capture all colony entries and exits. This setup enables the precise tracking of individual colony attendance patterns over their entire breeding cycle (see ref.^139^ and supplementary material F in ref.^141^ for a description of the system, as well as similar systems in Adélie^55,77,142^ and other penguins species^141,143–145^). The RFID record produced by this system was used to determine individual trip durations between the 2010 and 2024 breeding seasons (see trip duration section below).

Each season, this electronic monitoring was complemented by daily observations of a subset of nests spread over the study colony (n = 0 to 60 depending on the season), starting in 2012. We identified breeding partners on their nest using either hand-held RFID readers or repeated visual observations coupled with the automatic RFID system. We monitored both the breeding outcome and phenology (e.g., hatching and crèching dates) of each of these nests throughout the breeding season. Because nests were monitored from a distance using binoculars to minimize disturbance, the hatching date could not be measured precisely for all nests. Nests for which hatching dates could not be measured with a precision of ± 4 days (i.e. more than 4 days between the last observation of an egg and the first observation of a chick) were thus excluded from the analyses. Crèching dates were observed with a 1 -day precision. For a given nest, hatching dates were taken as the hatching of the first egg, and crèching dates as the first time the brood was seen unattended by parents. Finally, we conducted weekly photo counts of both adults and chicks present in the study colony from the 2011 season onwards. These counts were used to determine the reproductive success of the study colony (chicks fledged per breeding pair), which closely represented the broader archipelago population (r = 0.99, 2011 -2017, ref.^139^, see also^146^).

### Focus on the chick-guard stage

The chick-guard stage of the Adélie penguin starts at hatching and ends when chicks are left unattended at the colony while both parents forage simultaneously (crèching stage). We decided to focus our analyses on the chick-guard stage of the Adélie penguin breeding cycle for three reasons. First, foraging parents are most constrained in time and space during chick-guard. Compared to incubation and crèching, they forage over a smaller area^67^ and provision chicks more frequently^51,69,147^. These constraints on parental effort during chick-guard are further illustrated by the body mass loss incurred by breeders, whereas this pattern is typically reversed during incubation and crèching ^105,106^. Second, chicks have a higher probability of mortality during chick-guard compared to crèche^98,99^, making this stage critical for chick survival and reproductive success. Finally, focusing on chick-guard facilitates comparisons of foraging trips duration with other study sites, as data is more abundant for chick-guard than for crèching, in part because of the practical difficulties associated with retrieving electronic data loggers during crèching (another common methodology for acquiring trip duration data)^85,148,149^.

In Adélie penguins, the phenology of reproductive stages varies among years and populations ^51,150^. In Adélie Land, peak hatching and crèching typically occur in late December and mid-to-late January, respectively^151^. Here, we defined the chick-guard stage on an annual basis according to season-specific median hatching and crèching dates. Nest-level hatching dates were obtained from the RFID record by adding 31 days (i.e. the duration of incubation minus the average two days taken by the female to leave the colony after laying^51^ to the departure date of females for their first incubation foraging trip (average annual n = 77 ± 41 nests). Crèching dates were obtained from both daily observations at the colony and the RFID record, with crèching defined as the first time two known partners were away from the colony (average annual n = 25 ± 12 nests). Both hatching and crèching dates derived from the RFID record were validated using the visual observation data for the subset of nests for which both were available. As the field observations of crèching dates started in 2013, crèching phenology was unavailable in 2010, 2011, and 2012. For those years, we defined the median crèching date as the median hatching date plus the average duration of brooding in other years (25 days).

### Trip duration, sea ice, and individual parameters

Individual trip durations were extracted from the RFID record and defined as the duration (hours, h) between colony departure and return. We followed the data cleansing protocol of Bardon et al.^141^ to process missed detections (e.g., individuals detected with only one of the two directional antennas). Foraging trip data were also filtered to exclude trips shorter than 6 h (likely non-foraging^69,152^) and trips exceeding 5 days, as they likely reflect late incubation trips of late breeders, or unresolvable missing RFID detections. We ensured that we focused on trips from individuals engaged in chick -guarding activities by only using data from successful breeders, or from failed breeders prior to their breeding failure. To identify successful breeders, we used the *RFIDeep* workflow^141^, which identifies breeding success from individual RFID records with high accuracy (> 95%). For failed breeders, we retained only individuals for which the daily observation protocol (see above) allowed us to determine the date of breeding failure with a precision of ± 2 days or less.

We considered two sea ice variables to investigate how penguin foraging effort responded to sea ice variability and associated food availability. First, we assessed the binary presence/absence of landfast ice (sea ice anchored to the coastline, also called landfast ice) in front of the breeding colony during the chick-guard stage. This was achieved by visual inspection of MODIS and VIIRS satellite imagery using NASA Worldview (https://worldview.earthdata.nasa.gov). Second, we measured the sea ice concentration (SIC) in the chick-guard foraging area for each trip. A single SIC value per trip was calculated by averaging daily concentrations from one day pre-departure to one day post-return. Adélie penguins’ foraging area during chick-guard at our study site was defined using previously published tracking data^67,78,153^ and ranged from 65.5°S to 66.7°S and from 138°E to 141°E (Fig. S1). This spatial window is also consistent with recent findings that January SIC in that area was related to reproductive success for the Pointe Géologie archipelago Adélie penguin population^96^. Trip specific SIC was calculated using daily SIC data from Advanced Microwave Scanning Radiometer AMSR-E and AMSR-2 (6.25 × 6.25 km grid datasets), downloaded from the University of Bremen website (https://data.seaice.uni-bremen.de/amsr2/asi_daygrid_swath/) for all breeding seasons except 2012. AMSR data was unavailable for 2012 due to satellite replacement, and we instead used data from the National Snow and Ice Data Center^154^ (https://nsidc.org/data/g02202/versions/4, 25 × 25 km grid, in that season, as SIC from both datasets were strongly correlated for the 2009-2024 period (R^2^ = 0.96, β = 0.96 ± 0.005). The possible range for sea ice concentration values is between 0% (open water), and 100% (total ice cover).

In addition to landfast ice presence and SIC variability, we also included sex in our trip duration models, since female Adélie penguins usually forage for longer^76,84,86,155^, travel farther offshore^67^, and target different food sources^84,156^ compared to males. Sex assignment was done by analyzing sex-specific colony attendance patterns using RFIDeep^141^. Sex was determined by calculating the average percentage of support across multiple breeding seasons. Individuals for which the average percentage of support was below 75% (i.e. individuals with too little stereotyped colony-attendance patterns to be sexed confidently) were excluded from subsequent analyses (mean support for sex-assignation in the final dataset = 89%).

### Mean trip duration as an indicator of reproductive success

While our first hypothesis dealt with the effects of sea ice on foraging effort at the trip-scale, our second hypothesis focused on linking colony-scale measures of foraging effort to reproductive success. We tested our prediction that increased trip duration during chick-guard leads to lower reproductive success at two spatial scales. First, we averaged trip durations for each year at our Adélie Land study site and investigated the relationship between mean trip duration and reproductive success using both our entire dataset (n = 15 years) and a dataset only containing years when landfast ice was absent from the front of the colony. In both cases, we modelled reproductive success as a function of mean trip duration calculated over either all individuals, males, or females, and compared the three models using Akaike Information Criterion (AIC). Asreproductivesuccessdata for ourstudy colony was unavailable in 2010, we extracted it at the population level from ref.^96^ using PlotDigitizer (https://plotdigitizer.com/).

Second, we surveyed the literature using Google Scholar with the keywords ‘Adélie penguin’ and ‘trip duration’. For each identified study with trip duration data, we then looked for concurrent data on reproductive success, either from the same study or from a different study (see supplementary text). We extracted the reported trip duration data only when concurrent reproductive success data was available (i.e. same colony and same season). As there were slight differences among the 14 identified studies in how trip duration data was reported, we used our own data to ensure comparability of methods (see supplementary text for details).

### Statistical analyses

#### Trip duration and sea ice

We used Generalized Linear Mixed Models (GLMM) and Generalized Additive Mixed Models (GAMM) as implemented in the {*gamm*} function of R package{*mgcv*}^157^ to investigatelinear and non-linear relationships between trip duration and SIC. We fitted linear and quadratic GLMM as well as GAMMs, and compared these models to assess the support for non-linearity in the response of trip duration to SIC, using an AIC-based model selection framework^158^. As a conservative way to avoid model overfitting and to optimize biological interpretation, the maximum number of basis functions used for smoothing the relationship between trip duration and SIC in GAMMs was set to *k* = 4. In all models, we used a Gamma error distribution with a log link function and included individual ID as a random effect to account for individual-level repeated measurements. Additional fixed effects of sex, as well as the interaction of sex with SIC were then included, and their support was investigated using AIC. For GAMMs, we employed a tensor product smoother (t2)^159^ to allow sex-specific smoothing of the relationship between trip duration and SIC (i.e. interaction)^160^. Data from each landfast ice scenario (present/absent) was analyzed separately because there were large differences in both sample size and range of SIC values covered between scenarios.

### Reproductive success and mean trip duration: study site

To model the relationship between colony-scale mean trip duration and reproductive success (number of chicks fledged per pair) for our study site, we first averaged all trip duration values by year, without accounting for repeated measurements at the individual level (i.e. a slightly different approach to that of Emmerson and Southwell^76^ but similar to Ainley et al.^161^. We then fitted Generalized Linear Models (GLM) with a log link function as implemented in package {*glmmTMB*}^162^ with reproductive success and mean trip duration as response and explanatory variables, respectively. We fitted three models, first using annual mean trip duration for all individuals, then for males and females separately. Reproductive success was modeled using a Tweedie distribution to accommodate the presence of 0 (2 values out of 15) and the lower bound at 0^163^. We weighted observations using the inverse standard error of each annual trip duration mean, thus giving greater weight to more precise estimates of mean trip duration (i.e. measured over larger sample sizes, Table S1). We assessed the validity of the unweighted models by checking residuals for dispersion, uniformity, and outliers using package {*DHARMa*}^164^. We used a variance-based pseudo-R^2^ (squared correlation coefficient between predicted and observed data) to provide an indication of the predictive performance of our models.

### Reproductive success and mean trip duration: across colonies

We used the same approach to upscale the analysis using data from 9 other colonies. We fit Tweedie GLMM with a colony random effect to account for possible differences in mean reproductive success among colonies. We compared a linear and a quadratic model using AIC to test for non-linear effect of mean trip duration on reproductive success. Because standard errors for each annual mean trip duration were only available for <30% of the data entries, we weighted observations using a log-transformation of the number of trips used to calculate each annual mean (supplementary text). As mean trip duration estimates made over a larger number of trips are likely to be more precise (although asymptotically) ^165^, this allowed to put more weight on more precise estimates. Model fit was again assessed using the package{*DHARMa*}. Because initial checks revealed significant zero-inflation, we incorporated a zero-inflation component dependent on the explanatory variable in the Tweedie GLMMs. We used package {*performance*} to compute conditional and marginal R^2^ ^166,167^.

All statistical analyses were performed using R version 4.4.2^168^. Mean values are presented ± SE unless specified otherwise.

## Acknowledgements

We are deeply grateful to all the present and former members of Project 137, including the member of the MIBE team at IPHC in Strasbourg for their expertise and support, Yvon Le Maho (former PI of 137-ECOPHY), and all the wintering and summering field teams since the inception of this project in the field in 2005. We would also like to thank all the members of the missions in Dumont D’Urville since then, and the French Polar Institute-IPEV logistics team in Dumont d’Urville for their important and continuous support in the field. This study was approved by the French Polar Environmental Committee and authorizations for handling animals and accessing breeding sites were delivered by the Terres Australes et Antarctiques Françaises (TAAF).

## Data availability

Data and R scripts will be made available on Figshare upon acceptance of the manuscript.

## Authors contributions

Author contributions follow the CRediT guidelines. In each section, authors are listed by order of appearance in the author list.

Conceptualization: TB, NL, CLB

Methodology: TB, AH, MB, RC, TR, NL, CLB

Software: TB, GB

Validation: TB, NL, CLB

Formal analysis: TB, GB, NL, CLB

Investigation: TB, AH, MB, PB, RC, TR, DPZ, NL, CLB

Resources: CLB

Data Curation: TB, GB, CT, LL, CLB

Writing – original draft preparation: TB, NL, CLB

Writing – review and editing: All authors Visualization: TB

Supervision: NL, CLB Project administration: CLB

Funding acquisition: DPZ, NL, CLB

## Supplementary text

### Circumpolar analysis of the effect of trip duration on reproductive success using published data

To upscale our analysis relating chick-guard trip duration to reproductive success from our study to several populations around the species range, we surveyed the literature using Google Scholar with the keywords ‘Adélie penguin’ and ‘trip duration’. For each studyfor which trip duration data was available, we then looked for concurrent data on reproductive success, either from the same study or from a different study (see Supplementary Table S5). We extracted the reported annual trip duration summary statistics only when the reproductivesuccessdatawas available for samecolony and for the same season. Overall, we obtained 14 different studies to use in our circumpolar analysis.

Because the reporting of trip duration data varied among these 14 studies, we used our own dataset to compare the different methodologies and ensure comparability between our approach and those used by other authors. The differences between our approaches and those found in the literature can be summarized in three main themes:

### 1. Use of a different summary statistic

In one study (Watanabe et al. 2020), the summary statistics used to report annual trip duration was the median. In contrast, and to remain consistent with most other studies, we used the mean. Using our own data to compare mean and median trip duration, we found that *annual medians* were highly correlated to *annual means* (linear model: β = 0.98 ± 0.03, p < 0.001, R^2^ = 0.99).

### 2. Use of a different method to compute annual mean trip durations (means of means vs. grand mean)

#### Mean of means-per-period

In two studies, mean trip durations were reported separately for four (Ainley et al., 2004) to six (Saenz et al., 2020) periods of 5-6 days spanning the entire chick-rearing period. In one study (Ainley et al., 2006), mean trip duration were reported daily. In all three cases, we considered only periods falling during the chick-guard stage and averaged these period-specific means in order to obtain a single annual mean trip duration value. Using our own dataset, we compared this method (i.e. *mean of means-per-period*) to using the *grand mean* over all annual trips. Both methods yielded comparable estimates of annual mean trip duration, with the *mean of means-per-period* being a strong predictor of the *grand mean* both when using 6 equal-sized and successive subsamples within season (linear model: β = 0.99 ± 0.01, p < 0.001, R^2^ = 0.99) and when using daily means (linear model: β = 0.99 ± 0.02, p < 0.001, R^2^ = 0.99).

#### Mean of means-per-individual

In one study (Emmerson et al., 2015), individual means were first calculated for each individual (yielding a single mean trip duration value per individual per year) before performing the averaging of individual means, as recommended by Southwell et al. (2006). In contrast, and to ensure direct comparability with most other studies, we calculated mean trip duration using all trips directly (i.e. *grand mean*, without prior within-individual averaging). Using our own dataset, we compared both methods. We found that mean trip duration calculated as the *mean of means-per-individual* was highly comparable to the *grand mean* (linear model: β = 0.95 ± 0.02, p < 0.001, R^2^ = 0.99).

#### Mean of means-per-sex

In three studies (Clarke et al., 1998; Emmerson et al., 2015; Machado-Gaye et al., 2025), mean trip durations were reported separately for males and females. We averaged these sex-specific estimates in order to obtain a single annual mean trip duration value to use in our comparative analysis. Using our own dataset, we compared this method (i.e. *mean of means-per-sex*) to using the *grand mean* over all trips. Both methods yielded comparable estimates of annual mean trip duration, with the *mean of means-per-sex* being a strong predictor of the *grand mean* (linear model: β = 1.03 ± 0.02, p < 0.001, R^2^ = 0.99).

### 3. Data not restricted to the chick-guard stage

In three studies (Jennings et al., 2021; Lescroël et al., 2020; Watanabe et al., 2020), annual trip duration metrics were reported for the entire chick-rearing season (i.e. over both the brooding stage and part of the crèching stage). These studies accounted for 7 out of the 64 total data entries. Because we could not isolate trip duration for the guard-stage (i.e. as we did for (Ainley et al., 2004, 2006; Saenz et al., 2020; Watters et al., 2020), we used these estimates directly in our comparative analysis. Using a version of our own dataset that also included trip duration data during crèching (n = 25,304 additional foraging trips), we compared annual mean trip duration calculated only for the chick-guard stage to annual mean trip duration calculated for the entire chick-rearing period (chick-guard and crèching). We found that both values were acceptably comparable (linear model: β = 1.33 ± 0.13, p < 0.001, R^2^ = 0.88).

When the number of trips used to compute mean trip duration was not reported by the authors, we used the minimal possible value (e.g., number of individuals for which trip duration data is reported, neglecting the possibility that an individual contributes more than one trip to the dataset).

Overall, despite some minor difference in the way trip duration data was reported across studies, we are confident that our approach is reasonable and should not impact the overall model, for which the predicted error is larger than the among-studies differences highlighted here.

---

**Figure S1:**
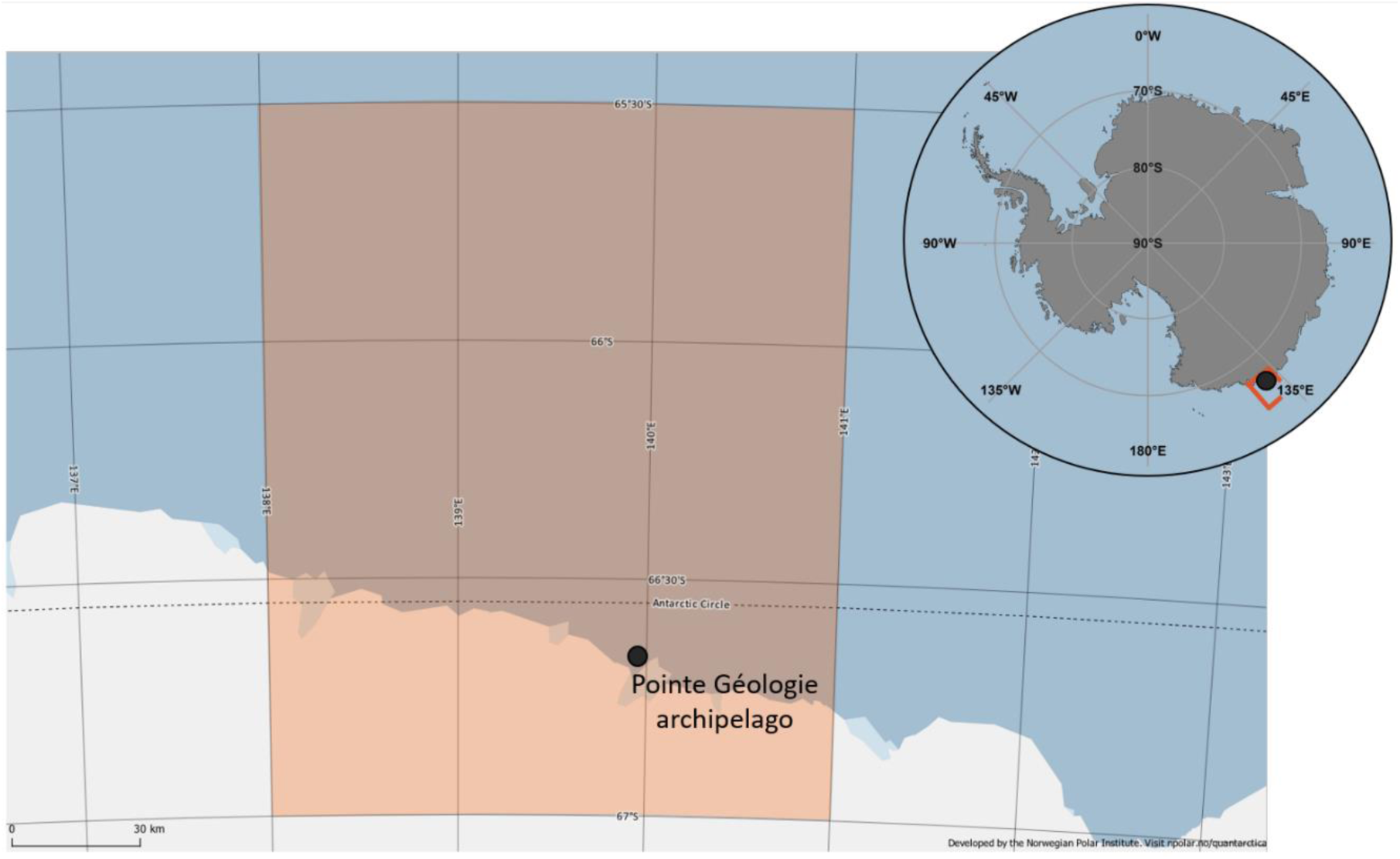
**Foraging area (orange) of chick-guarding Adélie penguins from the Pointe Géologie archipelago**, based on tracking data from Widmann et al. (2015), Michelot et al (2021), and Ropert-Coudert et al. (2018). The main map was produced using Quantarctica (Matsuoka et al. 2021) (https://www.npolar.no/quantarctica/).

**Figure S2:**
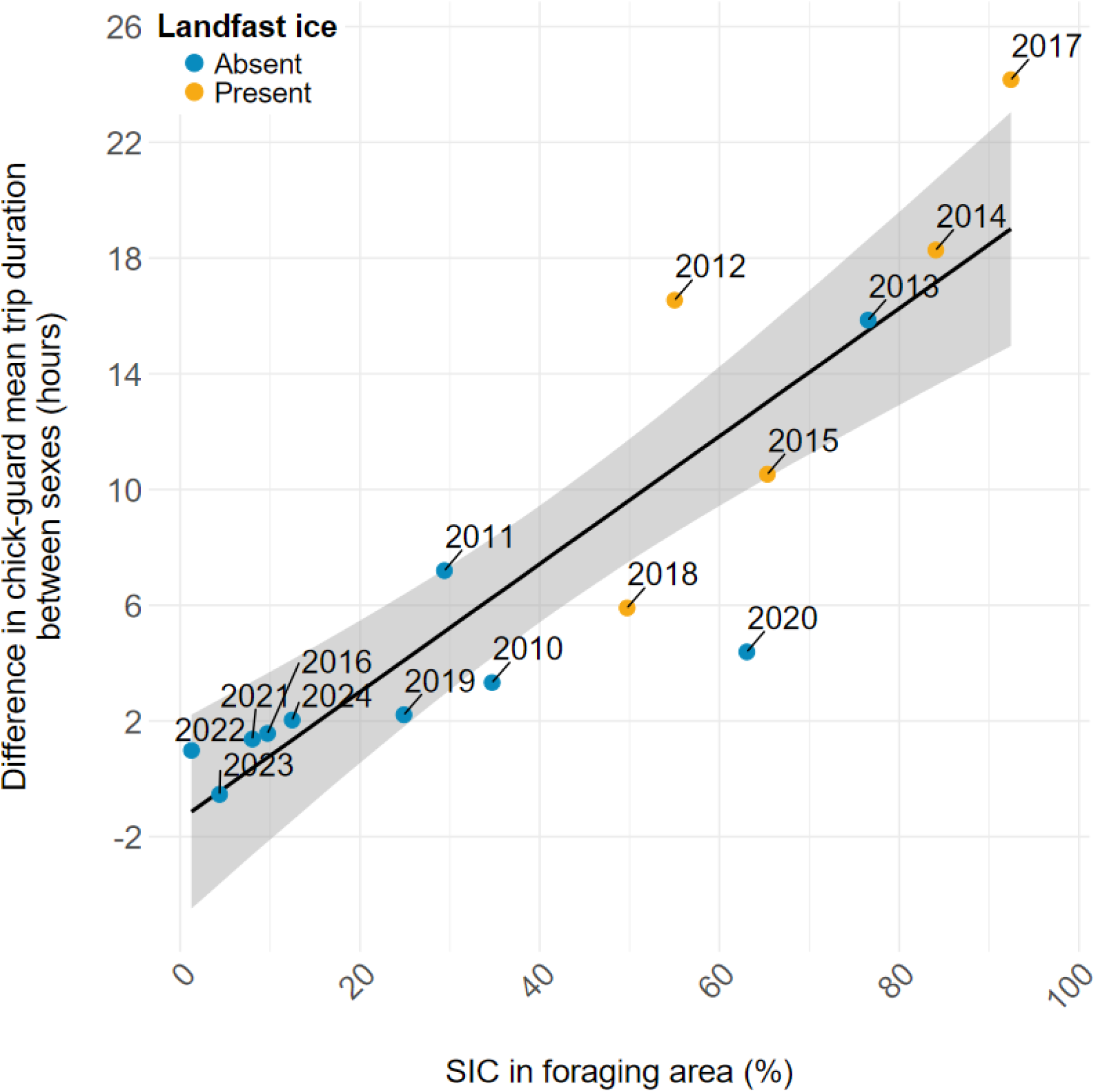
Relationship between the male-female difference in mean trip duration and Sea Ice Concentration (SIC) for chick-guarding Adélie penguin (*Pygoscelis adeliae*) breeding on Île des Pétrels, Pointe Géologie archipelago, Adélie Land, Antarctica (2010-2024, Linear Model, β = 0.22 ± 0.03, p < 0.001, R^2^ = 0.77). The landfast ice categorization indicates whether or not landfast ice was present in front of the breeding colony during chick-rearing, precluding direct access to open water. Higher values of the trip duration difference (y-axis) indicate longer trips by females compared to males.

**Figure S3:**
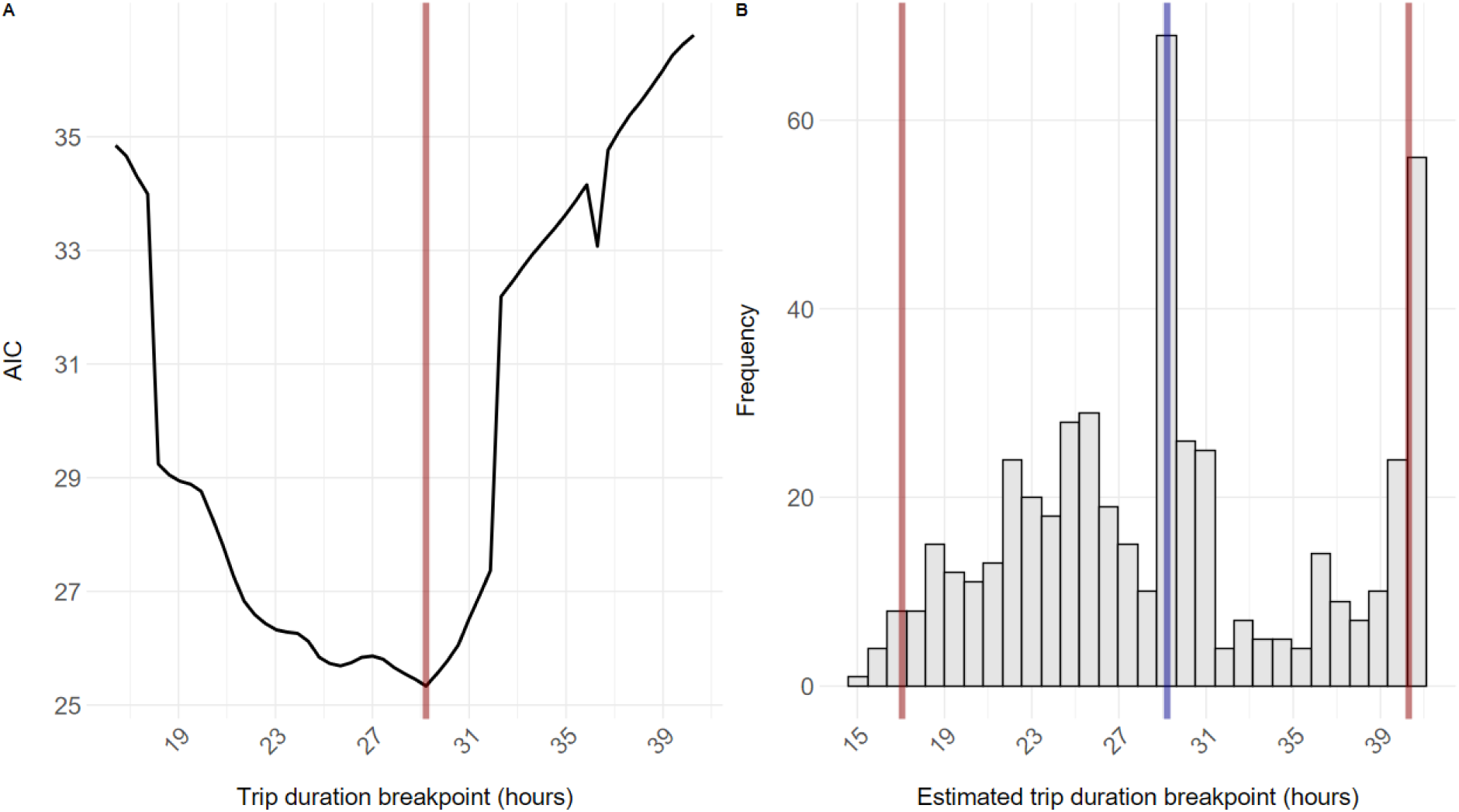
**A) Akaike Information Criterion (AIC) profile of the piecewise regression analysis examining the non-linear relationship between trip duration and reproductive success of Adélie penguins** (*Pygoscelis adeliae*) presented in Fig. 4, main text. The red vertical line denotes the most supported breakpoint (29.2 hours). **B) Distribution of bootstrapped breakpoint estimates** with the median (blue vertical line) and the 2.5th and 97.5th percentiles (red vertical lines). We conducted the piecewise regression analysis *a posteriori* to identify the trip duration threshold at which reproductive success begins to change at a different rate with increasing trip duration. Building on model 4 in Table S4, we evaluated 70 candidate breakpoints equally spaced between 10 and 40 hours of trip duration (i.e. ∼0.43 hours increment). We discarded models with less than 10 datapoints on either side of the breakpoint. We compared the remaining models using AIC to identify the most strongly supported breakpoint.To estimatea 95% confidenceinterval around this breakpoint we conducted a nonparametric bootstrap procedure by resampling the full dataset with replacement 500 times. For each bootstrapped sample, we re-estimated the optimal breakpoint by fitting a series of segmented Tweedie GLMMs across 70 candidate breakpoints, as explained above. The breakpoint associated with the lowest AIC was retained for each sample. We then derived the 95% bootstrap confidence interval calculated the 2.5^th^ and 97.5^th^ percentiles of the resulting breakpoint distribution to derive a 95% bootstrap confidence interval.

**Table S1:** Sample sizes and summary statistics of the sex-specific trip duration data. used to link trip duration during chick-guard to sea ice conditions and reproductive success for Adélie penguin (*Pygoscelis adeliae*) breeding on Île des Pétrels, Pointe Géologie archipelago, Adélie Land, Antarctica. N = *sample size*.

| Breeding season | N trips | N trips ♂ | N trips ♀ | N individuals ♂ | N individuals ♀ | N individuals | Sex ratio trips<br>(males prop.) | Sex ratio individuals<br>(males prop.) | Mean trip<br>duration ± SE | ♀ mean trip<br>duration ± SE | ♂ mean trip<br>duration ± SE |
| --- | --- | --- | --- | --- | --- | --- | --- | --- | --- | --- | --- |
| 2010 | 295 | 215 | 80 | 20 | 8 | 28 | 72.9 | 71.4 | 29.2 ± 0.6 | 31.7 ± 1.3 | 28.3 ± 0.7 |
| 2011 | 316 | 220 | 96 | 19 | 9 | 28 | 69.6 | 67.9 | 27.3 ± 0.7 | 32.3 ± 1.2 | 25.1 ± 0.8 |
| 2012 | 87 | 49 | 38 | 8 | 8 | 16 | 56.3 | 50 | 45.4 ± 2.1 | 54.8 ± 3.3 | 38.2 ± 2.1 |
| 2013 | 473 | 254 | 219 | 37 | 34 | 71 | 53.7 | 52.1 | 39.4 ± 1 | 48 ± 1.4 | 32.1 ± 1.2 |
| 2014 | 28 | 10 | 18 | 5 | 10 | 15 | 35.7 | 33.3 | 72.2 ± 4.4 | 78.8 ± 5.5 | 60.5 ± 5.7 |
| 2015 | 274 | 125 | 149 | 24 | 30 | 54 | 45.6 | 44.4 | 53 ± 1.4 | 57.8 ± 1.9 | 47.3 ± 2 |
| 2016 | 5297 | 2284 | 3013 | 111 | 147 | 258 | 43.1 | 43 | 17.4 ± 0.1 | 18.1 ± 0.1 | 16.5 ± 0.1 |
| 2017 | 35 | 20 | 15 | 10 | 9 | 19 | 57.1 | 52.6 | 34 ± 3.5 | 47.8 ± 5.9 | 23.7 ± 2.4 |
| 2018 | 1197 | 609 | 588 | 73 | 71 | 144 | 50.9 | 50.7 | 41.3 ± 0.5 | 44.3 ± 0.7 | 38.4 ± 0.6 |
| 2019 | 2433 | 1174 | 1259 | 96 | 99 | 195 | 48.3 | 49.2 | 23.1 ± 0.2 | 24.1 ± 0.2 | 21.9 ± 0.3 |
| 2020 | 2122 | 1028 | 1094 | 111 | 116 | 227 | 48.4 | 48.9 | 29 ± 0.2 | 31.1 ± 0.3 | 26.8 ± 0.3 |
| 2021 | 4255 | 2052 | 2203 | 110 | 116 | 226 | 48.2 | 48.7 | 18.9 ± 0.1 | 19.6 ± 0.2 | 18.2 ± 0.1 |
| 2022 | 2899 | 1486 | 1413 | 111 | 103 | 214 | 51.3 | 51.9 | 21.6 ± 0.1 | 22.1 ± 0.2 | 21.1 ± 0.1 |
| 2023 | 1381 | 650 | 731 | 51 | 51 | 102 | 47.1 | 50 | 24.2 ± 0.2 | 23.9 ± 0.2 | 24.4 ± 0.4 |
| 2024 | 2367 | 1148 | 1219 | 88 | 94 | 182 | 48.5 | 48.4 | 23.6 ± 0.2 | 24.6 ± 0.2 | 22.6 ± 0.2 |
| All (2010-2024) | 23459 | 11324 | 12135 | 260 | 305 | 565 | 48.3 | 46 | 23.4 ± 0.1 | 24.4 ± 0.1 | 22.4 ± 0.1 |

**Table S2:**
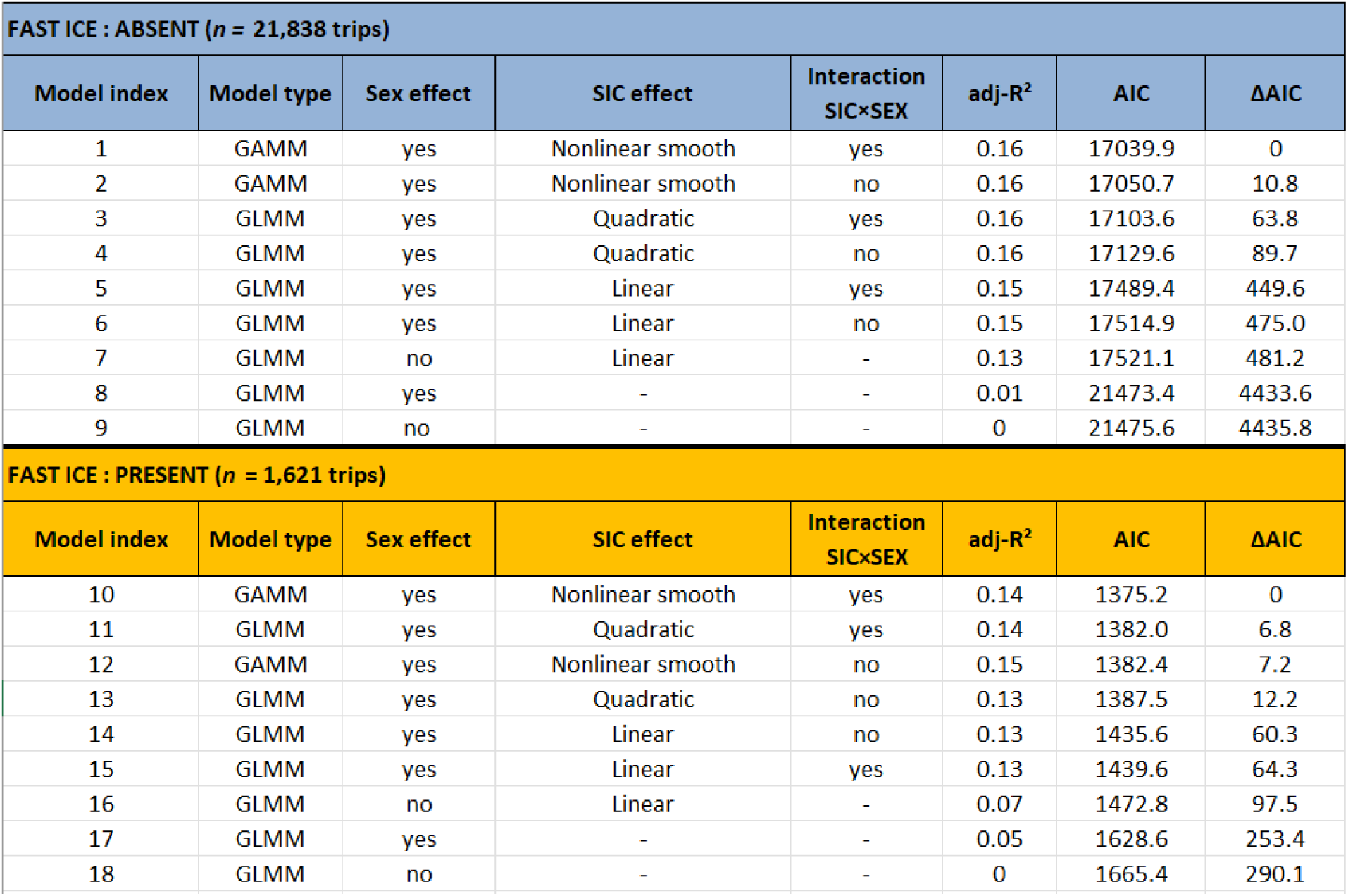
Model selection table for the effects of sea ice concentrations (SIC) and sex on the trip duration of chick-rearing Adélie penguins (*Pygoscelis adeliae*) during the guard stage in two scenarios of access to open water for breeding penguins: direct access to open water is either possible (*fast ice absent*) or precluded by the presence of solid landfast ice in front of the breeding colony (*fast ice present*). All models used trip duration as the response variable, included a random individual effect to account for repeated measurements on the same individuals, and used a Gamma error distribution with a log link. Trip duration datawas collected at Île des Pétrels, Pointe Géologie archipelago, Adélie Land, Antarctica. Abbreviations: GAMM: Generalized Additive Mixed Model; GLMM: Generalized Linear Mixed Model; adj-R^2^: percentage of variance explained by the fixed effect component of the mode; AIC: Akaike Information Criterion; ΔAIC: AIC difference between candidate model and model with the lowest AIC within the model set.

**Table S3:** Summary of models relating mean trip duration during chick-guard to reproductive success for Adélie penguin (*Pygoscelis adeliae*) at Pointe Géologie archipelago, Adélie Land, Antarctica. The effect of trip duration on reproductive success was modeled using Generalized Linear Models (GLM) with a Tweedie distribution and a log link. Pseudo R^2^ were calculated based on the percentage of observed variance in the data explained by the model. AIC: Akaike Information Criterion; ΔAIC: AIC difference between candidate model and model with the lowest AIC within the model set.

| Dataset | Mean trip duration<br>calculated over | Tweedie dispersion<br>parameter | Effect size | SE effect size | P value | Pseudo R<sup>2</sup> | AIC | ΔAIC |
| --- | --- | --- | --- | --- | --- | --- | --- | --- |
| All seasons (n=15) | All individuals | 0.046 | -0.044 | 0.006 | <0.001 | 0.73 | -22.17 | 0 |
| All seasons (n=15) | Females | 0.049 | -0.038 | 0.006 | <0.001 | 0.78 | -5.88 | 16.29 |
| All seasons (n=15) | Males | 0.052 | -0.049 | 0.008 | <0.001 | 0.57 | 3.83 | 26 |
| Seasons without landfast ice (n=10) | All individuals | 0.039 | -0.035 | 0.007 | <0.001 | 0.48 | -15.1 | 0 |
| Seasons without landfast ice (n=10) | Females | 0.046 | -0.03 | 0.008 | <0.001 | 0.48 | -9.71 | 5.39 |
| Seasons without landfast ice (n=10) | Males | 0.042 | -0.044 | 0.010 | <0.001 | 0.46 | -9.51 | 5.59 |

**Table S4:** Summary of models investigating linear and non-linear effects of foraging trip duration (annual mean) on reproductive success for chick-rearing Adélie penguins (*Pygoscelis adeliae*). All models included a random population effect (n = 10 populations, Fig. 4, Table S5) and a fixed effect of trip duration. Abbreviations: AIC: Akaike Information Criterion. ΔAIC: AIC difference between candidate model and model with the lowest AIC within the model set.

| Model index | Model | Fixed effect | Random effect | Fixed effect shape | Weighting | Tweedie dispersion parameter | Conditional R<sup>2</sup> | Marginal R<sup>2</sup> | AIC | ΔAIC |
| --- | --- | --- | --- | --- | --- | --- | --- | --- | --- | --- |
| 1 | Reproductive success ~ Trip Duration + (1\|Colony) | Mean trip duration | Colony | Quadratic | log(Ntrips) | 0.049 | 0.87 | 0.74 | -39.95 | 0 |
| 2 | Reproductive success ~ Trip Duration + (1\|Colony) | Mean trip duration | Colony | Linear | log(Ntrips) | 0.049 | 0.83 | 0.56 | -6.9 | 33.05 |
| Model index | Model | Fixed effect | Random effect | Fixed effect shape | Weighting | Tweedie dispersion parameter | Conditional R<sup>2</sup> | Marginal R<sup>2</sup> | AIC | ΔAIC |
| 3 | Reproductive success ~ Trip Duration + (1\|Colony) | Mean trip duration | Colony | Quadratic | none | 0.049 | 0.83 | 0.75 | 26.48 | 0 |
| 4 | Reproductive success ~ Trip Duration + (1\|Colony) | Mean trip duration | Colony | Linear | none | 0.065 | 0.65 | 0.49 | 36.89 | 10.41 |

**Table S5:** Literature review for concurrent data on Adélie penguin (*Pygoscelis adeliae*) foraging trip duration and reproductive success. Full size table with extracted data and references available at: https://figshare.com/s/ef3fe5b9ce965fc6ca13

